# Population Structure Analysis of Globally Diverse Bull Genomes

**DOI:** 10.1101/059139

**Authors:** Neo Christopher Chung, Joanna Szyda, Magdalena Fra̧szczak, the 1000 Bull Genomes Project

## Abstract

Since domestication, population bottlenecks, breed formation, and selective breeding have radically shaped the genealogy and genetics of *Bos taurus*. In turn, characterization of population structure among globally diverse bull genomes enables detailed assessment of genetic resources and origins. By analyzing 432 unrelated bull genomes from 13 breeds and 16 countries, we demonstrate genetic diversity and structural complexity among the global bull population. Importantly, we relaxed a strong assumption of discrete or admixed population, by adapting latent variable models for individual-specific allele frequencies that directly capture a wide range of complex structure from genome-wide genotypes. We identified a highly complex population structure that defies the conventional hypothesis based on discrete membership and contributes to pervasive genetic differentiation in bull genomes. As measured by magnitude of differentiation, selection pressure on SNPs within genes is substantially greater than that on intergenic regions. Additionally, broad regions of chromosome 6 harboring largest genetic differentiation suggest positive selection underlying population structure. We carried out gene set analysis using SNP annotations to identify enriched functional categories such as energy-related processes and multiple development stages. Our comprehensive analysis of bull population structure can support genetic management strategies that capture structural complexity and promote sustainable genetic breadth.

## Introduction

*Bos taurus* (cattle) has long experienced selection for high quality milk and meat production. To maintain and encourage genetic diversity, it is important to characterize the population structure of bulls around the world. Inferring population structure and genetic differentiation play an increasingly important role in conservation efforts, genealogy, and selection programs. In this study, we have analyzed a large number of whole genome sequences of *Bos taurus* bulls from 13 breeds, representing 16 countries, to characterize population structure and genetic diversity.

Recognizing the importance of cattle genome diversity in genome-wide association studies, genomic predictions, and optimal breeding, there have been substantial efforts to obtain genome-wide genotypes of multiple breeds in diverse geographical locations^1–3^. The 1000 Bull Genomes Consortium has successfully collaborated with institutions from more than 20 countries to collect 1577 whole genome sequences (as of version 5). This international collection of diverse genomes can be regarded as representative of genetic diversity within bulls and thus enables systematic analysis of population genomics. Although the structural complexity of cattle has previously been studied based on limited genome profiles or genetic markers, focusing on regions and breeds^4–7^, a population genetic study involving a large and diverse collection of whole genome sequences has not been performed.

Moreover,most studies assumed discrete structure among representatives of a studied population, as defined by self-identified breeds. Recent studies using genome-based tree, admixture models, and other techniques demonstrate far greater structural complexity^1,2,7^, but direct estimation and utilization of continuous population structure have been challenging. Logistic factor analysis (LFA) uses recently developed probabilistic models of individual allele frequencies underlying genotypes that are appropriate for a wide range of population structures (e.g., discrete, continuous, or admixture)^8^. Building on principal component analysis (PCA), LFA provides a non-parametric estimation method tailored to genotype data. By modeling each single nucleotide polymorphism (SNP) by the population structure estimated by logistic factors (LFs), genetic differentiation can be directly tested and inferred.

Applying latent variable probabilistic models, we analyzed 432 unrelated *Bos taurus* genomes from 13 breeds and 16 countries, as part of the 1000 Bull Genomes Project^2^. This study provides detailed assessment of population structure among a diverse panel of whole genome sequences (> 3.9 million SNPs per bull). We identified pervasive genetic differentiation as suggested by domestication and selection. Through incorporating gene set analyses with genomic features, evolutionary pressure on genetic variation is investigated. Additionally, we present an interactive visualization, which enables exploration of underlying population structure by logistic factors. This study represents one of the first studies in population genomics where potentially inaccurate breeds (or other self-referential subpopulation labels) are intentionally left unused.

## Results

In the 1000 Bull Genomes Project dataset, there were *n* = 432 unrelated *Bos taurus* samples with average sequencing coverage > 5 (Figure 1). These bulls represent 13 different breeds; namely, Angus, Brown Swiss, Charolais, Gelbvieh, Holstein, Jersey, Limousin, Montbeliard, Normandy, Piedmont, European Red Dairy, Holstein, Red & White, and Simmental/Fleckvieh. Defined by the official animal identification, our samples came from Australia, Austria, Canada, Denmark, Finland, France, Germany, Italy, Netherlands, New Zealand, Norway, Spain, Sweden, Switzerland, United Kingdom, and United States (Figure 2). Among these genomes, there are *m* = 3,967,995 single nucleotide polymorphisms (SNPs) with no missing values and minor allele frequencies ¿ 0.05 (Supplementary Fig. 2). To explore structural complexity, whole genome sequences of *n* = 432 selected samples were hierarchically clustered using Manhattan distances (Figure 3, colored by 13 different breeds). It is evident that official breed codes (or countries of origin) do not necessarily represent the genetic diversity among bulls represented by SNPs.

The dimension of the population structure in logistic factor analysis was set at *d* =7, as estimated by the VSS algorithm and the scree plot of decreasing eigenvalues (Supplementary Fig. 3). The estimated logistic factors demonstrate the genetic continuum, reflecting shared origins of genetics and overlapping goals of breeding programs since domestication (Figure 4). At the same time, the logistic factor 4 displays a clear distinction of Brown Swiss (from Switzerland, Germany, France, and Italy) and projection of logistic factors allows straightforward visual identification of clusters. We enable interactive exploration of this population structure by creating an online app visualizing logistic factors according to user-specified parameters (nnnn.shinyapps.io/bullstructure/).

**Figure 1.**
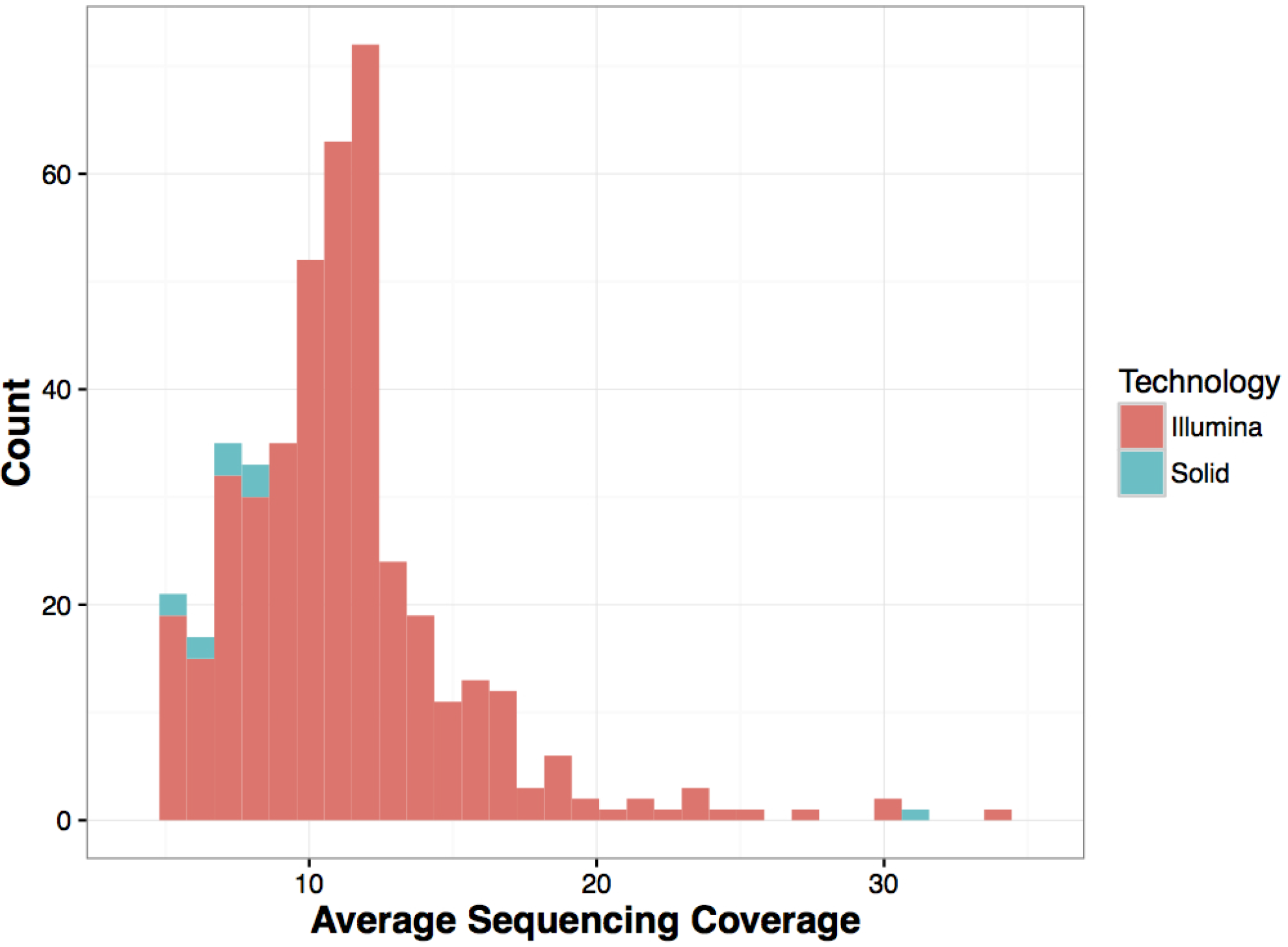
Average sequencing coverage of 432 bull samples.

**Figure 2.**
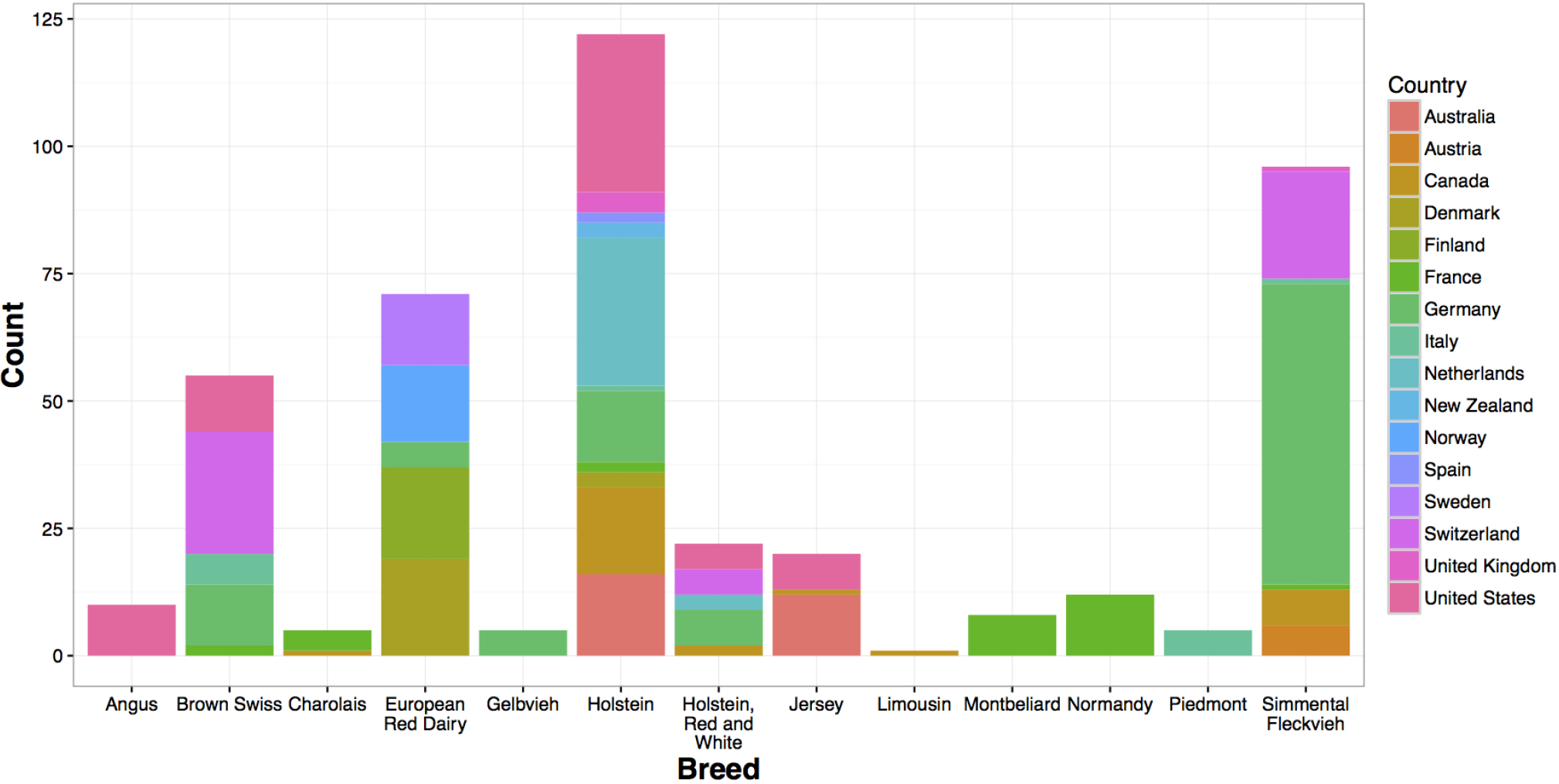
Bar plot of cattle breeds, with a number of samples colored by countries of origin.

**Figure 3.**
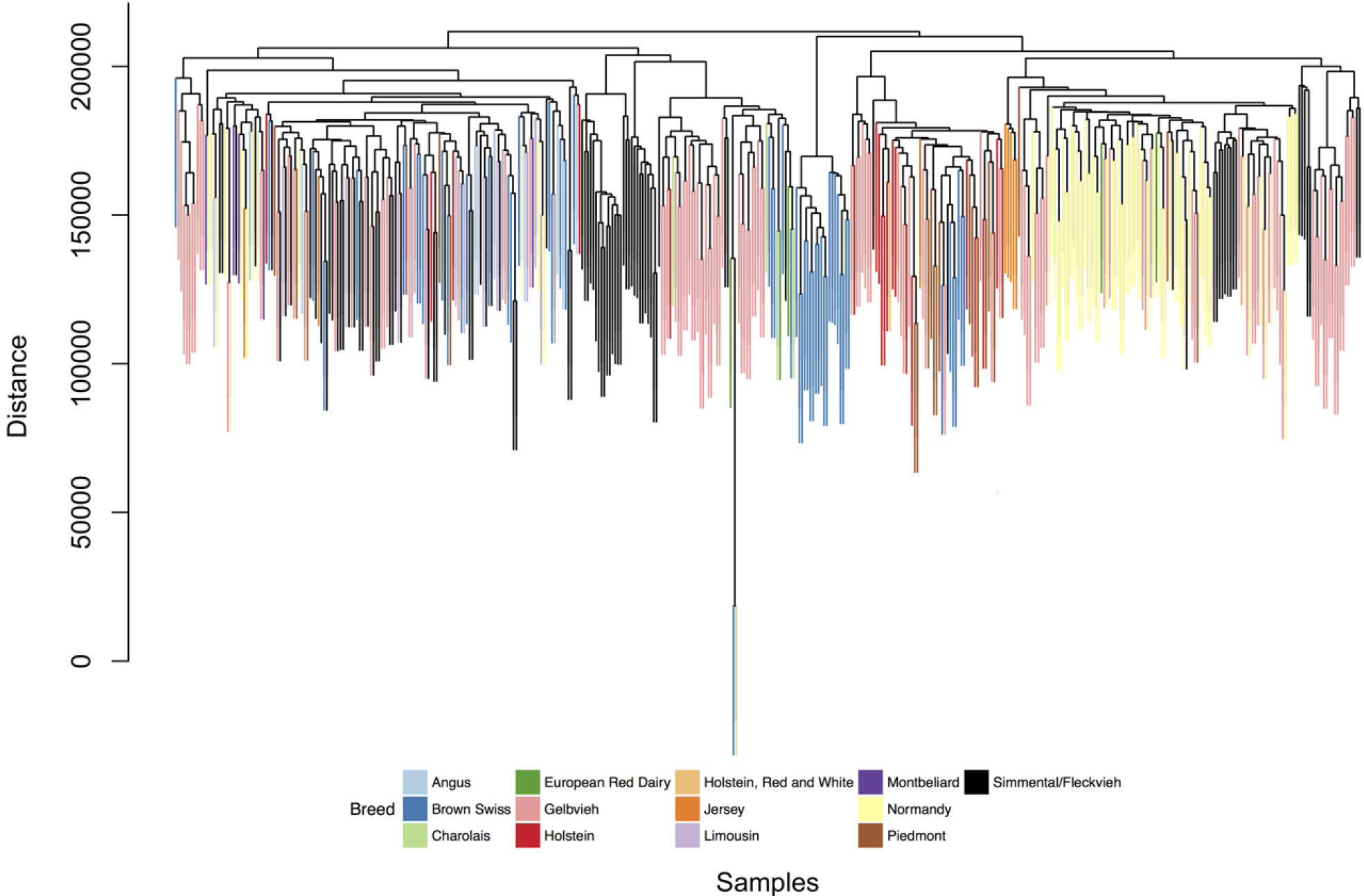
Hierarchical clustering of 432 bull genomes. Genome-wide SNPs are clustered using Manhattan distances and samples are colored by breeds.

**Figure 4.**
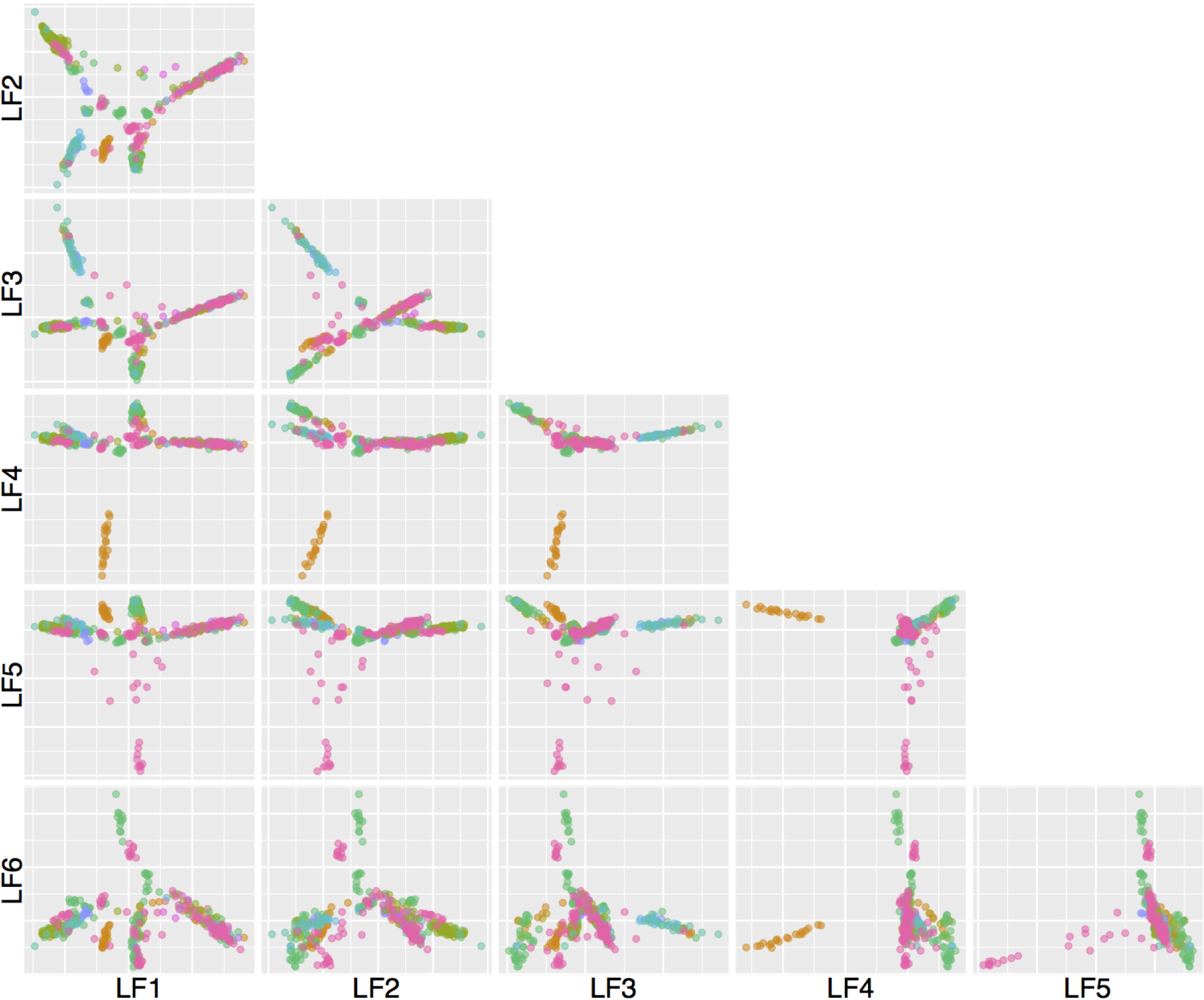
Scatterplots of logistic factors (LFs). All pairs of 6 LFs are plotted, excluding the intercept term. Data points corresponding to 432 bull genomes are colored by 13 breeds. Interactive visualization available at nnnn.shinyapps.io/bullstructure/.

We discovered diverse and pervasive genetic differentiation with respect to the population structure of bulls. When applying the resampling-based jackstraw method to test association between SNPs and logistic factors, we observed that the vast majority of SNPs are statistically significantly differentiated (an estimated proportion of null SNPs 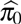 of 6.2%). A proportion of variation explained by *d* = 7 LFs for each SNP is approximated by McFadden’s pseudo *R*^2^. We found that the median and mean values of McFadden’s pseudo *R*^2^ are 0.070 and 0.087, respectively (Figure 5). The chromosome 6 contained substantially more SNPs with high *R*^2^ than other chromosomes; it harbors 166 (39.0%) out of 426 SNPs with *R*^2^ > 0.6, as well as all 29 (100%) SNPs with *R*^2^ > 0.7. On the other hand, the X chromosome shows the least variation with respect to logistic factors, containing zero SNP with *R*^2^ > 0.5. The top 1000 genomic features that are associated with differentiated SNPs are shown in Supplementary Data 1. Note that we additionally conducted an independent robustness analysis with *d* = 22 logistic factors (as suggested by a cross-validation method), which confirm highly consistent genetic differentiation, with an overall *R*^2^ correlation of 0.93.

**Figure 5.**
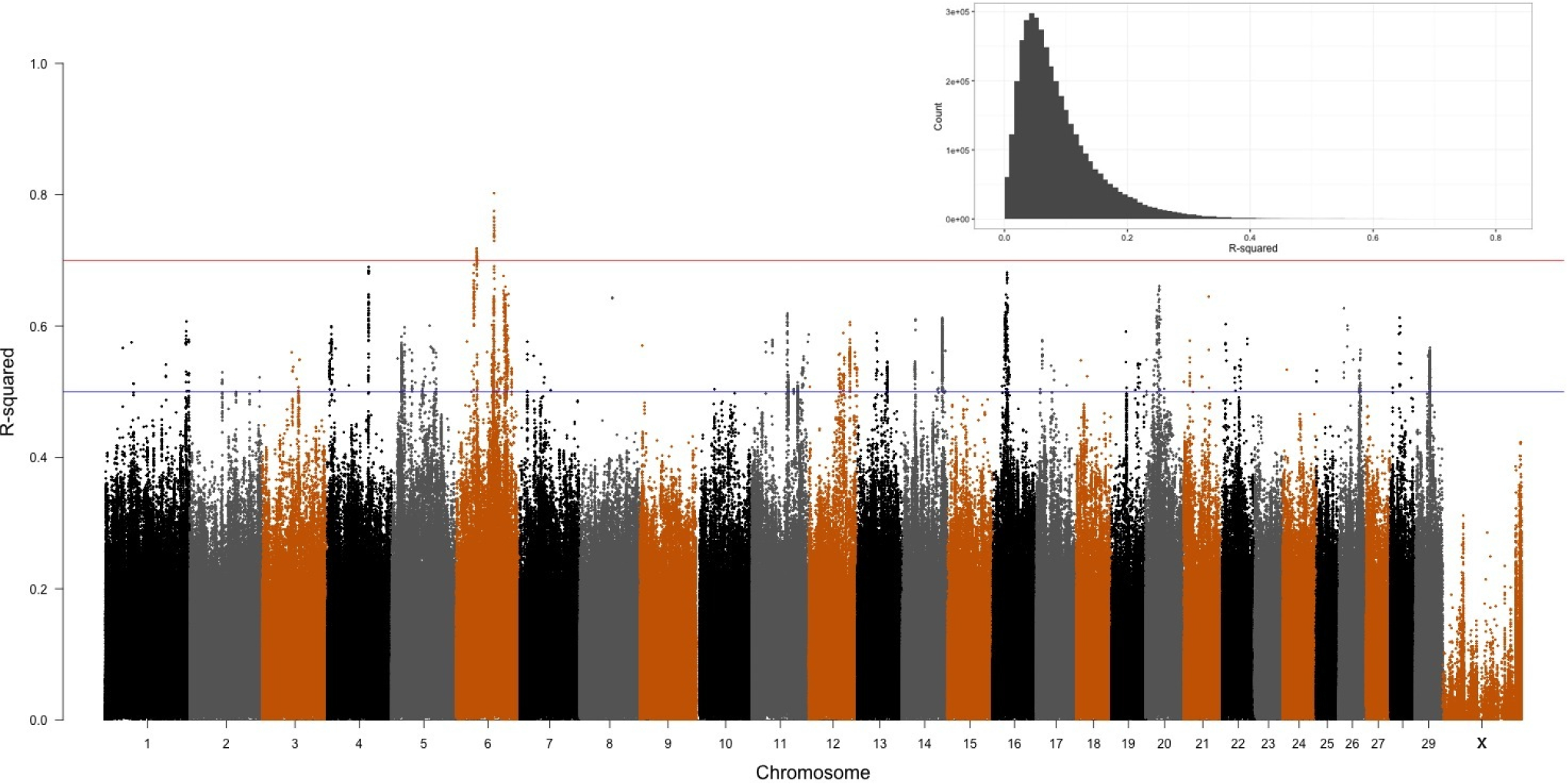
Genome-wide pseudo R^2^ measures with respect to logistic factors (LFs). The distribution is highly skewed towards 0, which leads to overplotting in a low range (see an insert for a genome-wide histogram). Overall, the median and mean are 0.070 and 0.087, respectively

Among SNPs with the highest *R*^2^ > 0.7, there exist two tight groups on chromosome 6; specifically 14 SNPs (13 within 50kbp of known genomic features) positioned between 71101370 and 71600122 and 15 SNPs (11 within 50kbp of known genomic features) positioned between 38482423 and 39140537. 83% of those most differentiated SNPs (20 out of 24 SNPs with known genomic features) are within or close to genes related to the selection sweep according to^9^. Among the first group, five SNPs fall within CHIC2 (ENSBTAG00000032660), while the closest features within 50kbp also include GSX2 (ENSBTAG00000045812), U6 spliceosomal RNA (ENSBTAG00000042948), and novel pseudogene (ENSBTAG00000004082). U6 spliceosomal RNA (ENSBTAG00000042948) and novel pseudogene (ENSBTAG00000004082) are known to be associated with milk protein percentage^10^. In the second group, the exact overlaps occur in FAM184B (ENSBTAG00000005932), LCORL (ENSBTAG00000046561), and NCAPG (ENSBTAG00000021582). LCORL encodes a transcription factor whose human ortholog is involved in spermatogenesis, whereas NCAPG is crucial in mitosis and meiosis. Expecting much granular investigation of such genomic features, the list of 396,800 SNPs at the top 90 percentile (*R*^2^ > .174) is available as Supplementary Data 2.

To better understand evolutionary and biological processes, we conducted gene set analyses using genomic annotations of SNPs. Firstly, we found that SNPs located within known genomic features have about 1.8% higher *R*^2^ measures than intergenic SNPs without annotations (MWW p-values 9.85 ⨯ 10^−106^). On the other hand, among intergenic SNPs, we found no significant correlation (p-value of 0.44) between SNP-feature distances and *R*^2^ measures (Supplementary Fig. 4). Secondly, among genic SNPs, *R*^2^ measures corresponding to SNPs within exons are slightly higher than those within introns by 0.27% with a MWW p-value 3.89 ⨯ 10^−29^. Start/stop codons and 3’/5’ UTR do not exhibit statistically significant difference from other genic SNPs. Lastly, we used 338 genes that are closest to SNPs with *R*^2^ > 0.5 in the `DAVID` functional annotation tools. We found a total of 34 enriched annotation clusters, of which 11 clusters with enrichment scores > 0.5 are shown in (Table 1). Biological processes and functions related to calcium-binding domain (cluster 1 and 9) and iron containing hemeproteins related to ATP (cluster 3 and 6) exhibit strong enrichment, potentially reflecting causes of population structure. Notably, we observed functional clusters for sexual, respiratory, and embryonic developments (cluster 5, 7, and 10, respectively).

**Table 1.** Enriched functional clusters, for genes associated with *R*^2^ >0.5

|  |  |  |  |  |
| --- | --- | --- | --- | --- |
| **Cluster 1** | **Enrichment Score: 1.405** | **Calcium-binding domain** |  |  |
| Category | Term | Count | % | PValue |
| INTERPRO | IPR018247:EF-HAND 1 | 6 | 2.098 | 0.035 |
| INTERPRO | IPR018249:EF-HAND 2 | 6 | 2.098 | 0.038 |
| INTERPRO | IPR011992:EF-Hand type | 6 | 2.098 | 0.045 |
| **Cluster 2** | **Enrichment Score: 1.372** | **Cysteine-type activity** |  |  |
| Category | Term | Count | % | PValue |
| GOTERM_MF_FAT | GO:0004198~calcium-dependent cysteine-type endopeptidase activity | 3 | 1.049 | 0.011 |
| GOTERM_MF_FAT | GO:0008234~cysteine-type peptidase activity | 4 | 1.399 | 0.066 |
| GOTERM_MF_FAT | GO:0004197~cysteine-type endopeptidase activity | 3 | 1.049 | 0.106 |
| **Cluster 3** | **Enrichment Score: 0.897** | **Cytochrome** |  |  |
| Category | Term | Count | % | PValue |
| PIR_SUPERFAMILY | PIRSF000045:cytochrome P450 CYP2D6 | 3 | 1.049 | 0.013 |
| INTERPRO | IPR002401:Cytochrome P450, E-class, group I | 3 | 1.049 | 0.068 |
| INTERPRO | IPR017973:Cytochrome P450, C-terminal region | 3 | 1.049 | 0.080 |
| INTERPRO | IPR017972:Cytochrome P450, conserved site | 3 | 1.049 | 0.084 |
| SP_PIR_KEYWORDS | heme | 4 | 1.399 | 0.091 |
| INTERPRO | IPR001128:Cytochrome P450 | 3 | 1.049 | 0.107 |
| SP_PIR_KEYWORDS | Monooxygenase | 3 | 1.049 | 0.124 |
| COG_ONTOLOGY | Secondary metabolites biosynthesis, transport, and catabolism | 3 | 1.049 | 0.148 |
| GOTERM_MF_FAT | GO:0020037~heme binding | 4 | 1.399 | 0.159 |
| GOTERM_MF_FAT | GO:0046906~tetrapyrrole binding | 4 | 1.399 | 0.176 |
| GOTERM_MF_FAT | GO:0009055~electron carrier activity | 4 | 1.399 | 0.301 |
| SP_PIR_KEYWORDS | iron | 4 | 1.399 | 0.399 |
| GOTERM_MF_FAT | GO:0005506~iron ion binding | 4 | 1.399 | 0.614 |
| **Cluster 4** | **Enrichment Score: 0.860** | **Signaling** |  |  |
| Category | Term | Count | % | PValue |
| UP_SEQ_FEATURE | signal peptide | 19 | 6.643 | 0.048 |
| SP_PIR_KEYWORDS | signal | 19 | 6.643 | 0.111 |
| SP_PIR_KEYWORDS | glycoprotein | 16 | 5.594 | 0.492 |
| **Cluster 5** | **Enrichment Score: 0.833** | **Sexual development** |  |  |
| Category | Term | Count | % | PValue |
| GOTERM_BP_FAT | GO:0045137~development of primary sexual characteristics | 3 | 1.049 | 0.117 |
| GOTERM_BP_FAT | GO:0003006~reproductive developmental process | 4 | 1.399 | 0.151 |
| GOTERM_BP_FAT | GO:0007548~sex differentiation | 3 | 1.049 | 0.180 |
| **Cluster 6** | **Enrichment Score: 0.760** | **Ion binding** |  |  |
| Category | Term | Count | % | PValue |
| GOTERM_MF_FAT | GO:0043167~ion binding | 40 | 13.986 | 0.130 |
| GOTERM_MF_FAT | GO:0046872~metal ion binding | 38 | 13.287 | 0.190 |
| GOTERM_MF_FAT | GO:0043169~cation binding | 38 | 13.287 | 0.213 |
| **Cluster 7** | **Enrichment Score: 0.725** | **Respiratory development** |  |  |
| Category | Term | Count | % | PValue |
| GOTERM_BP_FAT | GO:0030324~lung development | 3 | 1.049 | 0.145 |
| GOTERM_BP_FAT | GO:0030323~respiratory tube development | 3 | 1.049 | 0.145 |
| GOTERM_BP_FAT | GO:0060541~respiratory system development | 3 | 1.049 | 0.150 |
| GOTERM_BP_FAT | GO:0035295~tube development | 3 | 1.049 | 0.400 |
| **Cluster 8** | **Enrichment Score: 0.723** | **Protease activity** |  |  |
| Category | Term | Count | % | PValue |
| GOTERM_MF_FAT | GO:0004175~endopeptidase activity | 8 | 2.797 | 0.129 |
| GOTERM_MF_FAT | GO:0070011~peptidase activity, acting on L-amino acid peptides | 9 | 3.147 | 0.190 |
| GOTERM_MF_FAT | GO:0008233~peptidase activity | 9 | 3.147 | 0.215 |
| GOTERM_BP_FAT | GO:0006508~proteolysis | 12 | 4.196 | 0.242 |
| **Cluster 9** | **Enrichment Score: 0.703** | **Calcium-binding domain** |  |  |
| Category | Term | Count | % | PValue |
| UP_SEQ_FEATURE | calcium-binding region:2 | 3 | 1.049 | 0.126 |
| INTERPRO | IPR002048:Calcium-binding EF-hand | 4 | 1.399 | 0.148 |
| UP_SEQ_FEATURE | calcium-binding region:1 | 3 | 1.049 | 0.157 |
| SMART | SM00054:EFh | 4 | 1.399 | 0.187 |
| UP_SEQ_FEATURE | domain:EF-hand 1 | 3 | 1.049 | 0.258 |
| UP_SEQ_FEATURE | domain:EF-hand 2 | 3 | 1.049 | 0.258 |
| INTERPRO | IPR018248:EF hand | 3 | 1.049 | 0.333 |
| **Cluster 10** | **Enrichment Score: 0.668** | **Embryonic development** |  |  |
| Category | Term | Count | % | PValue |
| GOTERM_BP_FAT | GO:0001824~blastocyst development | 3 | 1.049 | 0.082 |
| GOTERM_BP_FAT | GO:0001701~in utero embryonic development | 4 | 1.399 | 0.165 |
| GOTERM_BP_FAT | GO:0043009~chordate embryonic development | 4 | 1.399 | 0.397 |
| GOTERM_BP_FAT | GO:0009792~embryonic development ending in birth or egg hatching | 4 | 1.399 | 0.400 |
| **Cluster 11** | **Enrichment Score: 0.565** | **Cardiomyopathy** |  |  |
| Category | Term | Count | % | PValue |
| KEGG_PATHWAY | bta05412:Arrhythmogenic right ventricular cardiomyopathy (ARVC) | 3 | 1.049 | 0.240 |
| KEGG_PATHWAY | bta05410:Hypertrophic cardiomyopathy (HCM) | 3 | 1.049 | 0.277 |
| KEGG_PATHWAY | bta05414:Dilated cardiomyopathy | 3 | 1.049 | 0.304 |
| **Cluster 12** | **Enrichment Score: 0.519** | **Phosphorylation** |  |  |
| Category | Term | Count | % | PValue |
| GOTERM_MF_FAT | GO:0004672~protein kinase activity | 9 | 3.147 | 0.213 |
| GOTERM_BP_FAT | GO:0006468~protein amino acid phosphorylation | 9 | 3.147 | 0.291 |
| GOTERM_BP_FAT | GO:0016310~phosphorylation | 9 | 3.147 | 0.447 |

## Discussion

*Bos taurus* has played a crucial role in ancient and modern societies alike by providing agricultural support and essential nutrients. Accurate characterization of its population structure helps conservation of genetic resources and optimal selection programs, ensuring a healthy and sustainable cattle population. In this process, we can better infer the genetic and functional variation that underlies the population structure. Our study, using 432 whole genome sequences of unrelated *Bos taurus* samples, provides a comprehensive and rigorous assessment of population structure among diverse bulls.

Assumptions underlying population structure and its estimation methods based on genotypes have evolved to address growing genomics data in terms of complexity and scales^11–13^. Particularly, the contentious and ambiguous definition of breeds merely requires “certain distinguishable characteristics” and practically relies on self-referential breed registry for sire and dam^14^. Therefore, we have relaxed strong assumptions often used in population genomics, by employing latent variable probabilistic models^8^. In this study, we did not assume discrete or admixed populations. The present framework can nonetheless capture a broad range of arbitrarily complex structure including the aforementioned configurations. Based on observing a largely continuous genetic spectrum compared to breeds, we demonstrated that breeds do not account for structural complexity. We speculate that many cattle breeds, including presumed founders, are not as isolated or discrete as one would be led to believe. The population structure of our bull genomes in 7-dimensional logistic factors can be explored on our interactive visualization website.

When modeling SNPs with logistic factors in generalized linear models, we found widespread genetic differentiation due to population structure. This likely arose from a long history of breed formation and artificial selection, such that different national breeding programs may have caused weak and pervasive systematic variation. Despite being blind to breeds, the majority of the most differentiated SNPs in our study have been identified as under selection sweep. The chromosome 6 harboring a large proportion of highly differentiated SNPs has been suggested for strong selective sweep^1^, and may also be associated with calving ease and carcass weight^15,16^. Interestingly, given that the novel pseudogene (ENSBTAG00000004082), which has been known to be associated with calving performance^17^ and protein percentage^10^ is strongly associated with population structure, we suspect that it plays a crucial functional role in bull genomes. Overall, our genome-wide study of differentiation suggests stronger evolutionary pressure on genic regions. Enrichment analysis of genome annotations provides strong indications that functional groups related to energy production and development stages underlie the systematic variation in the panel of diverse bulls.

Pedigrees were used to remove 72.6% of bull samples related by progenitors, resulting in a panel of 432 genomes analyzed. However, undocumented kinship may potentially bias our population structure analysis, just as it does other methods that utilize breed and other subpopulation information. We advocate for stronger linkage between breeding programs and registries. The structural complexity among bull genomes discovered in this study can be used directly to identify genetic association with quantitative traits^18^. However, the 1000 Bull Genomes Consortium does not collect quantitative traits as its main goal is comprehensive identification of genomic variants. Lastly, the 1000 Bull Genomes Project, which is among the largest collections in this area of study, is still lacking samples from Asia, Africa, and South America.

This study paves a way to further our understanding of genetic diversity among modern cattle breeds. Our identification of systematic genetic differentiation may inform conservation efforts to preserve heritage breeds and maintain genetic diversity. Considering our flexible assumption about population structure and exclusive use of whole genome sequences, our highly differentiated SNPs, gene set analysis, and functional enrichment show how we can dispense of potentially inaccurate subpopulation labels in population genomics.

## Methods

### Bull Genomes

The 1000 Bull Genomes Project has collaborated with worldwide institutions to gather whole-genome sequences of diverse breeds. Its initial efforts have vastly expanded known single nucleotide polymorphisms (SNPs) and copy number variations (CNVs) in *Bos taurus^2^*. Currently, it covers 1577 bull samples as of version 5 released in 2015, among which 1507 and 70 bull genomes were sequenced with Illumina/Solexa and ABI SOLiD technology, respectively. For analysis of population structure, we selected unrelated bulls with average sequencing coverage greater than 5. Among sibs only one representative was selected randomly. SNP genotypes were identified prior to our study based on whole genome sequence data of bulls, using a multi-sample variant calling procedure. Polymorphisms with minor allele frequencies below 0.05 were removed from analyses. For processing whole-genome sequences, we used `vcftools v0.1.14`^19^, `BEDOPS v2.4.15`^20^, and `R v3.2.2`^21^.

### Statistical Analysis

To infer population structure directly from a genome-wide genotype matrix, we consider a probabilistic model of individual allele frequencies. In particular, by using logistic factor analysis^8^ that captures systematic variation of individual-specific allele frequencies arising from discrete or continuous sub-population, spatial variation, admixture, and other structures, we relax statistical assumptions imposed on bulls by its official breed and country code defined in the animal registration ID. While the statistical models and algorithms are extensively described in elsewhere^8^, we provide a brief overview of this approach here.

Consider a genotype matrix **Y** with *m* SNPs and *n* bulls. For each *y_ij_*, an individual-specific allele frequency for *i*^th^ SNP and *j^th^* bull is *f_ij_* ϵ [0,1]. This collection of parameters (a *m* ⨯ *n* **F** matrix) is transformed into real numbers via the logit function, which allows computation of the underlying latent structure. Overall, the statistical model considered is 
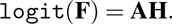
Then, the population structure is captured by d logistic factors (LFs) **H** which can be estimated by applying principal component analysis (PCA) to logit(**F**). Note that **A** is a matrix of coefficients in a logistic regression. The dimensions of logistic factors are estimated by comparing the observed correlation matrix to a series of hypothesized structures derived from selected variables of large loadings^22^. In the Very Simple Structure (VSS) algorithm, we considered *d* = 1,…, 100, while applying principal component analysis on the mean-centered genotypes (R package `psych`). Eigenvalues of *m*^−1^ **Y**^*T*^**Y** and percent variance explained by each component are visually inspected for the inflection point (e.g., elbow). For robustness analysis to confirm genetic differentiation, we alternatively used cross-validation approximations to choose *d*^23^.

To investigate genetic differentiation with respect to the population structure, we test association between *i*^th^ SNP y_i_ and estimate logistic factors Ĥ. We model SNPs with *d* logistic factors in a logistic regression (with a logit link function), where the deviance statistic compares the full (LFs) model *Y* ~ Ĥ and the null (intercept) model *Y* ~ 1^24^. We take account of the fact that the population structure is directly estimated from **Y** by utilizing the resampling-based jackstraw method^25^. For each of *B* iterations, the jackstraw method introduces a small number *s* ≪ *m* of permuted SNPs under a null model **Y** ~ 1 and computes *s* empirical null deviance statistics. P-values are calculated by ranking observed deviances with an empirical distribution of *B* ⨯ *s* deviances, as adapted from the resampling-based jackstraw approach^25^. This method based on a logistic regression is implemented in the jackstraw v1.1 package, freely available on the Comprehensive R Archive Network (cran.r-project.org/web/packages/jackstraw). A proportion of SNPs that are not associated with LFs (*π*_0_) is then estimated from *m* p-values.

To approximate how much of the variation in genotypes is explained by the population structure, we calculate McFadden’s pseudo *R*^2^ that is appropriate for a logistic regression^26^. This methodology is operationally similar to detecting genomic signatures with PCA^27^, although the difference arises from directly modeling categorical SNP data. For *i*^th^ SNP,
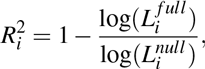
where 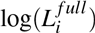 and 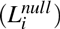 are maximum log-likelihoods of the full and null models, respectively. As this study only considers McFadden’s pseudo *R*^2^ in logistic regressions, we will henceforth refer to it as *R*^2^ when clear in context.

## Annotation and Enrichment

For genome annotation, we used the latest *Bos taurus* reference genome from the Center for Bioinformatics and Computational Biology, University of Maryland (downloaded from the NCBI server ftp://ftp.ncbi.nlm.nih.gov/, version UMD3.1.83).

When testing whether the distribution of McFadden’s pseudo *R*^2^ measures are significantly different according to feature types, we used the Mann-Whitney-Wilcoxon (MWW) test^28^. With a large sample size, a Normal approximation is used to compute MWW p-values. In particular, we investigated whether SNPs falling within genes may have a higher McFadden’s pseudo *R*^2^ than those in intergenic regions. Among SNPs with known feature assignments, MWW tests were used to infer if a particular feature type is associated with significantly higher *R*^2^ measures.

Lastly, because some of SNPs are in intergenic regions with no known annotations, we utilized the closest features function from `BEDOPS v2.4.15`^20^. Among the top genes with McFadden’s pseudo *R*^2^ > 0.5, we apply `DAVID v6.7` considering GO, KEGG pathways, InterPro, SwissProt Protein Information Resource, and other databases to identify enrichment of biological processes and functional pathways^29^. For intergenic SNPs, we searched the reference genome for the closest genes, which were used in `DAVID v6.7.` When clustering functional annotations, we set “Classification Stringency” to high.

## Acknowledgements

This work was supported by grant Polish National Science Centre (NCN) grant 2014 /13 / B / NZ9 / 02016. Part of data storage and computation were carried out at the Poznan Supercomputing and Networking Centre. N.C.C. was supported by the Leading National Research Center Programme 04/KNOW2/2014. The membership of the 1000 Bull Genomes Project are: Hans Rudolf Fries, Mogens SandϕLund, Bernt Guldbrandtsen, Didier Boichard, Paul Stothard, Roel Veerkamp, Michael Goddard, Curtis P Van Tassell, and Ben Hayes.

## Author contributions statement

N.C.C. conceived the study, analyzed data, wrote the first draft. M.F. contributed to editing the data. N.C.C. and J.S. revised the manuscript and contribute to the discussion.

## Additional information

N.C.C., M.F., J.S., and the 1000 Bull Genomes Project have no competing financial interest.

